# Chemical Genomics Language Model toward Reliable and Explainable Compound-Protein Interaction Exploration

**DOI:** 10.1101/2024.02.13.580100

**Authors:** Takuto Koyama, Hayato Tsumura, Ryunosuke Okita, Kimihiro Yamazaki, Aki Hasegawa, Keiko Imamura, Takashi Kato, Hiroaki Iwata, Ryosuke Kojima, Haruhisa Inoue, Shigeyuki Matsumoto, Yasushi Okuno

## Abstract

Deep learning-based compound-protein interaction (CPI) prediction models are promising in the field of molecular biology, particularly for facilitating the drug discovery process. In practical applications, CPI models should achieve a high generalization performance, quantify prediction confidence, and ensure explainability. Here, we propose ChemGLaM, a chemical genomics language model for reliable and explainable CPI predictions, by addressing these three crucial aspects. ChemGLaM integrates independently pre-trained chemical and protein language models through an interaction block with a cross-attention mechanism, achieving state-of-the-art performance in predicting novel CPIs. Incorporating uncertainty estimation and attention visualization enables ChemGLaM to enhance the success rate of virtual screening and to provide molecular insights into CPIs. Furthermore, we demonstrate its practical applicability by constructing a public database for large-scale CPI predictions and enabling drug/target exploration for candidate treatment of amyotrophic lateral sclerosis (ALS). ChemGLaM represents a significant step toward overcoming the challenges of AI-driven drug discovery and addressing unmet medical needs.

## Main

Identifying compound-protein interactions (CPIs) is crucial in molecular biology, particularly for facilitating the drug discovery process^1^. Exploring the vast chemical space to identify new hit compounds for novel targets is essential for drug development, particularly in areas with unmet medical needs^2^. Because the experimental validation of this vast chemical space is time-consuming and costly, computational methods provide a promising approach for predicting CPIs^3^. Deep learning (DL) is a powerful approach for CPI prediction, with many models adopting chemical genomics-based (structure-free) methods^4–6^.

Chemical genomics-based methods integrate chemical and genomic spaces using compound structures and protein sequences as inputs. The rapid advancement in DL algorithms, such as graph neural networks^5,7^, convolutional neural networks^8,9^, and transformer^4,10^ architectures, has enabled these models to efficiently learn the representations of two-dimensional compound structures and protein sequences from CPI data. In contrast to structure-based methods, such as molecular docking^11^ and molecular dynamics simulations, chemical genomics-based methods do not rely on the availability of accurate three-dimensional (3D) protein structures or require high computational costs, making large-scale CPI prediction feasible for target proteins^12^. Despite this progress, the practical applications of DL-based CPI prediction models are limited. Current CPI models substantially rely on the known data used in training, complicating the adaptation to novel compounds and proteins. While various CPI datasets have been used to build CPI prediction models^13–16^, they contain only tens of thousands to millions of samples, which is significantly smaller than the datasets used in computer vision (e.g., 14M images in ImageNet^17^) and natural language processing (e.g., 2.6T tokens in Llama 2^18^). Considering that the chemical genomics space combines the chemical (e.g., 119M compounds in PubChem) and protein (e.g., 300M sequences in UniRef) spaces, the current training data represents only a small portion of this vast chemogenomic space^19^. In addition to the limitations regarding the generalization performance for unknown CPIs, existing state-of-the-art models only provide prediction results and lack confidence scores and explainability. These aspects are crucial for preventing the wastage of development resources owing to incorrect predictions and for uncovering potential insights into the interaction mechanisms. A DL-based approach with all three aspects (high generalizability, confidence estimation, and explainability) may enhance the practical drug development process.

Here, we propose ChemGLaM, a chemical genomics language model designed to enhance the generalization performance, confidence, and explainability in CPI prediction. Our approach builds on the success of pre-trained language models, particularly chemical language models (CLMs) and protein language models (PLMs). These models mitigate data scarcity by learning meaningful representations from large-scale unlabelled datasets through self-supervised learning, facilitating adaptation to downstream tasks with limited data^20,21^. Because CPI outcomes depend on compound and protein properties, we integrated CLM and PLM using a multihead cross-attention mechanism. This enabled the model to capture relevant interactional features by leveraging the enriched representations learned by the CLM and PLM. Additionally, we applied an uncertainty estimation method, Evidential Deep Learning (EDL)^22^, to ChemGLaM to quantify the confidence of the model predictions.

Our contributions are as follows (Figure 1a):

(1) **Zero-shot CPI prediction**: We fine-tuned ChemGLaM for CPI prediction in a zero-shot setting, achieving state-of-the-art performance on benchmark datasets.
(2) **Uncertainty estimation**: ChemGLaM estimates prediction uncertainty, demonstrating its effectiveness in selecting reliable drug candidates through virtual screening.
(3) **Attention visualization**: We visualized the attention weights, providing explanatory insights into the mechanisms of CPI.

Furthermore, we demonstrated the impact on practical applications through the following analyses:

- **Open-source database publication**: We predicted all the CPIs between drugs and human proteins and published the results in an open-source database.
- **Drug screening for amyotrophic lateral sclerosis (ALS)**: Application of ChemGLaM to an experimental system using patient-induced pluripotent stem cells (iPSCs) showed the potential to identify new inhibitors and target proteins.

This study offers a scalable and efficient approach to improving CPI prediction accuracy and explainability, facilitating drug discovery.

**Figure 1:**
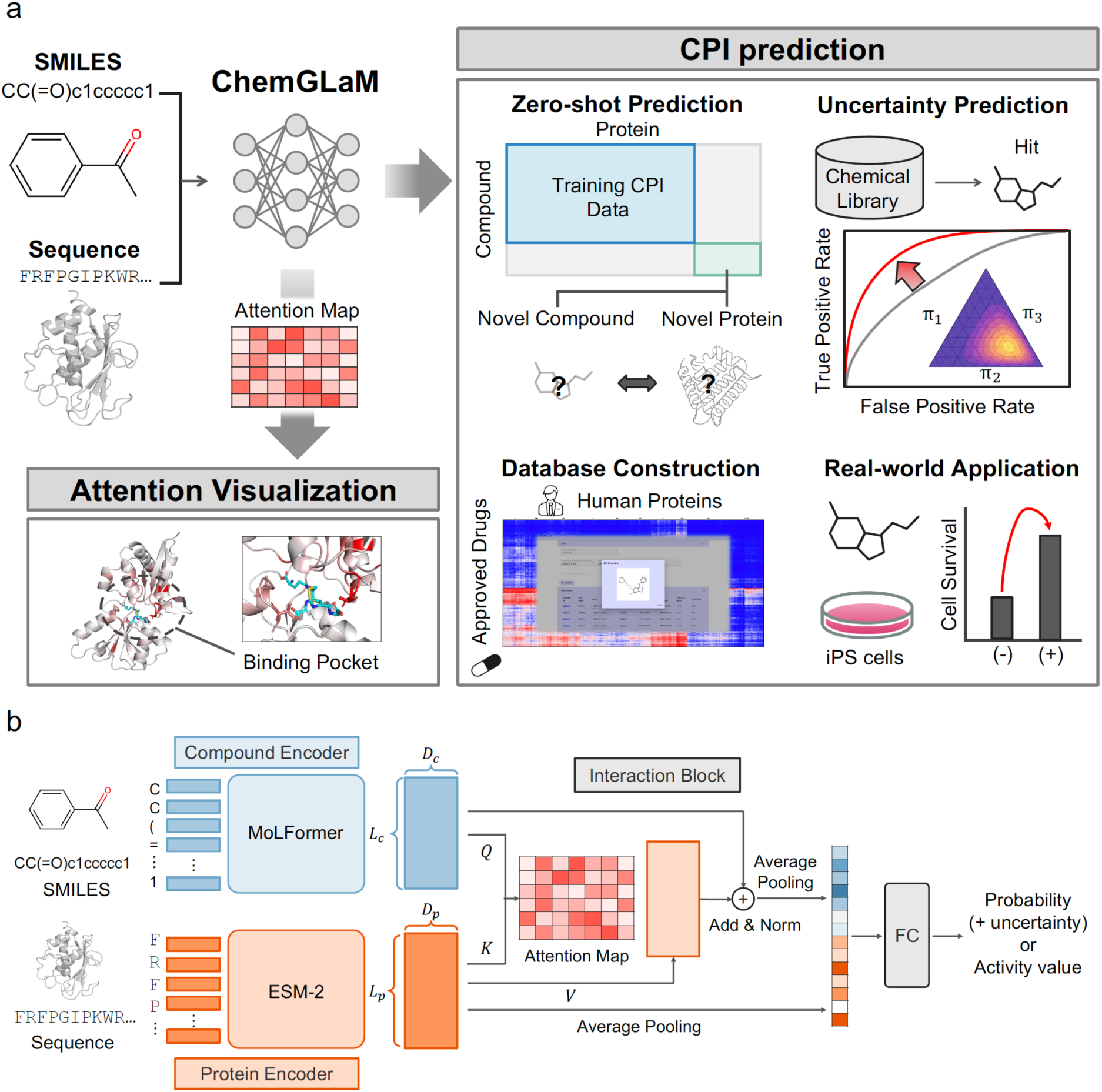
Schematic diagram of ChemGLaM. **a)** Overview of this study. **b)** ChemGLaM comprises three parts: Compound Encoder, Protein Encoder, and Interaction Block. The Compound Encoder is a chemical language model, MoLFormer, which processes a simplified molecular input line entry system (SMILES). The Protein Encoder is a protein language model, evolutionary scale modelling version 2 (ESM-2), which processes protein sequences. The Interaction Block connects the embeddings with cross-attention mechanisms (Q: query, K: key, V: value), and the output features are processed through fully connected (FC) layers. The notations and symbols used in the diagrams are defined in the Methods section.

## Results

### ChemGLaM architecture

ChemGLaM is an integrated model of PLM and CLM and comprises three parts: (1) a Compound Encoder, (2) a Protein Encoder, and (3) an Interaction Block (Figure 1b). The compound and protein encoders were derived from MoLFormer and evolutionary scale modelling version 2 (ESM-2), respectively, with pre-trained model parameters. Their embeddings were connected by an Interaction Block. Given that a compound interacts with a folded protein through regions that are distant in the primary sequence but spatially close to the 3D structure, the model should capture long-range relationships between compound atoms and protein residues for accurate CPI prediction. To learn the aspects of molecular interactions, a cross-attention mechanism was used as the Interaction Block to dynamically focus on different parts of the input data. Additionally, we implemented a function to estimate the uncertainty in CPI predictions using EDL. This function quantifies the confidence in ChemGLaM predictions by modelling the probability distributions over the predicted outcomes. These methods are detailed in the Methods section.

### ChemGLaM demonstrates state-of-the-art performance for unknown CPIs

To verify the generalization performance for unknown CPIs, we conducted model evaluations similar to zero-shot predictions (Supplementary Figure 1). We implemented 5-fold cross-validation on the test sets, excluding the compounds and proteins in the training datasets. BindingDB^15^ and Davis^13^ were used as datasets for classification, whereas regression tasks (binding-affinity predictions) comprised PDBbind^16^ and Metz^23^.

First, we compared the performance of the ChemGLaM model on the BindingDB dataset for zero-shot predictions with other existing CPI prediction models, hereafter referred to as baseline models (Figure 1a, Extended Data Table 1). ChemGLaM consistently demonstrated outstanding performance across multiple evaluation metrics. It achieved state-of-the-art performance with an area under the receiver operating characteristic curve (AUROC) of 0.829 (±0.042) and an accuracy of 0.754 (±0.039). This high performance indicates the effectiveness of large-scale pre-training in enhancing the generalization capabilities of ChemGLaM for predicting truly unknown CPIs. Similarly, the model excelled, exhibiting an area under the precision-recall curve (AUPRC) of 0.827 (±0.055) and an F1 score of 0.756 (±0.021), suggesting a high success rate of positive classifications, emphasising its overall efficiency in the dataset. In addition, ChemGLaM outperformed the baseline models on the Davis dataset, confirming its versatility (Figure 1b, Extended Data Table 1).

**Table 1:** Performance evaluation of ablation studies on ChemGLaM.

|  | BindingDB | Davis | PDBbind | Metz |
| --- | --- | --- | --- | --- |
| Model | AUROC | AUROC | Pearson | Pearson |
| ChemGLaM | **0.829 (0.042)** | **0.728 (0.047)** | 0.600 (0.026) | **0.544 (0.025)** |
| w/o pretraining MoL | 0.748 (0.037) | 0.731 (0.065) | **0.608 (0.036)** | 0.490 (0.044) |
| w/o MoLFormer | 0.673 (0.045) | 0.624 (0.014) | 0.423 (0.082) | 0.153 (0.040) |
| w/o ESM-2 | 0.827 (0.055) | 0.672 (0.066) | 0.553 (0.033) | 0.472 (0.045) |

AUROC: Area under the receiver operating characteristic curve.

Subsequently, we evaluated the performance of binding-affinity prediction for the PDBbind dataset, focusing on their performance in terms of root mean squared error (RMSE), Pearson correlation coefficient, Spearman correlation coefficient, and mean absolute error (MAE). Compared with those of the baseline models, ChemGLaM achieved an RMSE of 1.521 (±0.047), a high Pearson correlation of 0.600 (±0.026), a high Spearman correlation of 0.596 (±0.024), and a low MAE of 1.198 (±0.031) (Figure 1c, Extended Data Table 2). Notably, ChemGLaM predicts binding affinities more accurately than does the Vina scoring function (Spearman correlation coefficient of 0.511±0.037), a method based on molecular simulations. Conventional supervised approaches such as GraphDTA and TransformerCPI exhibit lower zero-shot performance than did Vina because of insufficient structural information and limited labelled data. ChemGLaM, which leverages large-scale pre-training, surpasses Vina even when relying solely on sequence information. In addition, we evaluated the performance of binding-affinity prediction for the Metz dataset and confirmed that ChemGLaM outperformed the baseline models across all metrics (Figure 1d, Extended Data Table 2).

These results demonstrate that ChemGLaM overcomes the limitations of supervised methods, considering truly unseen CPIs for classification and regression tasks, by combining CLM- and PLM-captured inherent representations. This generalizability highlights the potential of ChemGLaM for practical applications in drug discovery.

### ChemGLaM predicts distant CPIs with high generalizability

Although ChemGLaM achieved a state-of-the-art performance in zero-shot prediction, the test sets used in this study included compounds (or proteins) that were structurally (or sequentially) similar to those in the training set. To further understand the generalizability of ChemGLaM, we analysed its ability to predict distant pairs. Distant pairs refer to CPIs that have low compound or protein (or both) similarities to those of the training data. The similarities between compounds, proteins, and pairs were computed using Tanimoto similarity, sequence similarity (local alignment score), and their harmonic means (Methods and Figure 2e– g). Based on these similarities, we evenly divided the BindingDB test data into two groups: close (lower similarity) and distant (higher similarity) pairs. For close pairs, ChemGLaM maintained high predictive accuracy (AUROC of 0.819 (±0.083), 0.813 (±0.046), and 0.810 (±0.077) for Tanimoto, sequence, and CPI similarities, respectively), demonstrating a consistent advantage over baseline models reported earlier (Figure 2h–j). Notably, ChemGLaM also outperformed the baseline models on distant pairs, with AUROC of 0.831 (±0.035), 0.828 (±0.053), and 0.844 (±0.039) for Tanimoto, sequence, and CPI similarities, respectively. These results emphasise that ChemGLaM can be generalized for novel and challenging CPI pairs. We conducted the same evaluation for the Davis dataset and confirmed similar results (Supplementary Figure 2a–f).

**Figure 2:**
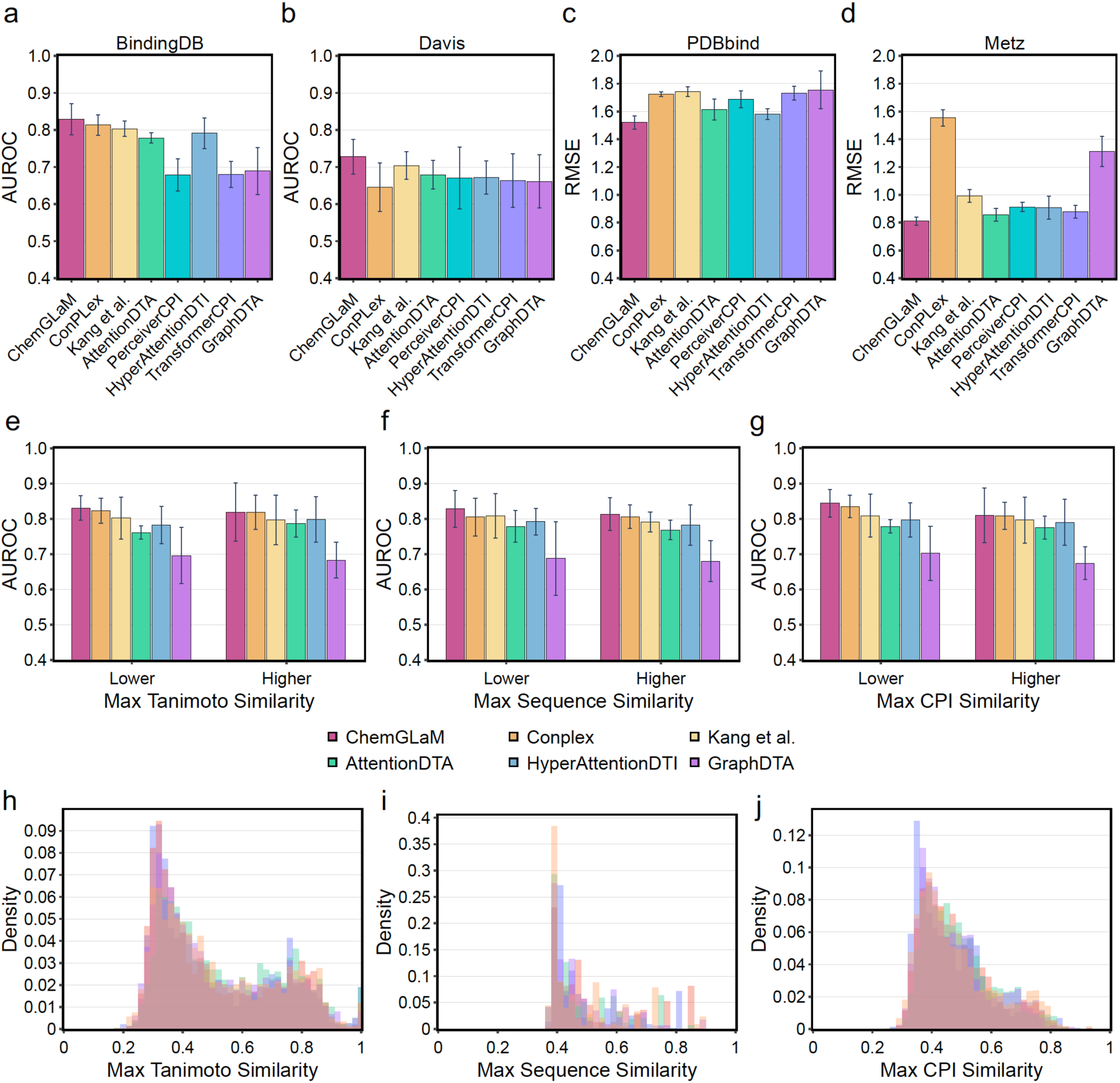
Performance evaluation of ChemGLaM in zero-shot prediction. **a-d)** Performance evaluation on BindingDB, Davis, PDBbind, and Metz datasets. **e–g)** The average area under the receiver operating characteristic curve (AUROC) scores of 5-fold cross-validation for the BindingDB dataset. The test set in each fold is divided into two groups based on the maximum similarities with the training set: close and distant pairs. The similarities are calculated via three approaches: Tanimoto, sequence, and compound-protein interaction (CPI) similarities. The CPI similarity is a harmonic mean of Tanimoto and sequence similarities. The bars on the left side represent the AUROC scores for the distant (lower similarity) pairs, and those on the right side represent the scores for the close (higher similarity) pairs. **h–j)** Distributions of maximum Tanimoto, sequence, and CPI similarities in five folds are represented. The distributions in each fold are indicated by different colours.

### ChemGLaM quantifies model uncertainty using evidential deep learning

Understanding confidence in a model’s prediction is essential for enhancing the success rate of virtual screening. Accordingly, we implemented methods within ChemGLaM to infer the model uncertainty using the EDL. In EDL, the neural network outputs positive values representing “evidence” for each class, which are treated as parameters for the Dirichlet Distribution. This approach allows the model to estimate uncertainty based on evidence (see the Methods section). We applied EDL to the BindingDB dataset and compared the uncertainty scores by stratifying a single test set based on three types of similarities with the training set: Tanimoto, sequence, and CPI (Figure 3a–c). Across all settings, test samples with high similarities exhibited low uncertainty scores, whereas the scores tended to deviate towards higher values as the similarity decreased. This indicates that the EDL implemented in ChemGLaM effectively detects out-of-domain data, allowing the appropriate interpretation of the prediction results by excluding them from the prediction targets. When samples with high uncertainty scores were excluded from the prediction target, ChemGLaM achieved a higher AUROC score (0.888) than did the entire test set (AUROC = 0.784) (Figure 3d). This result indicates that utilising uncertainty estimation improves the success rate of ChemGLaM-based virtual screening for practical drug discovery.

**Figure 3:**
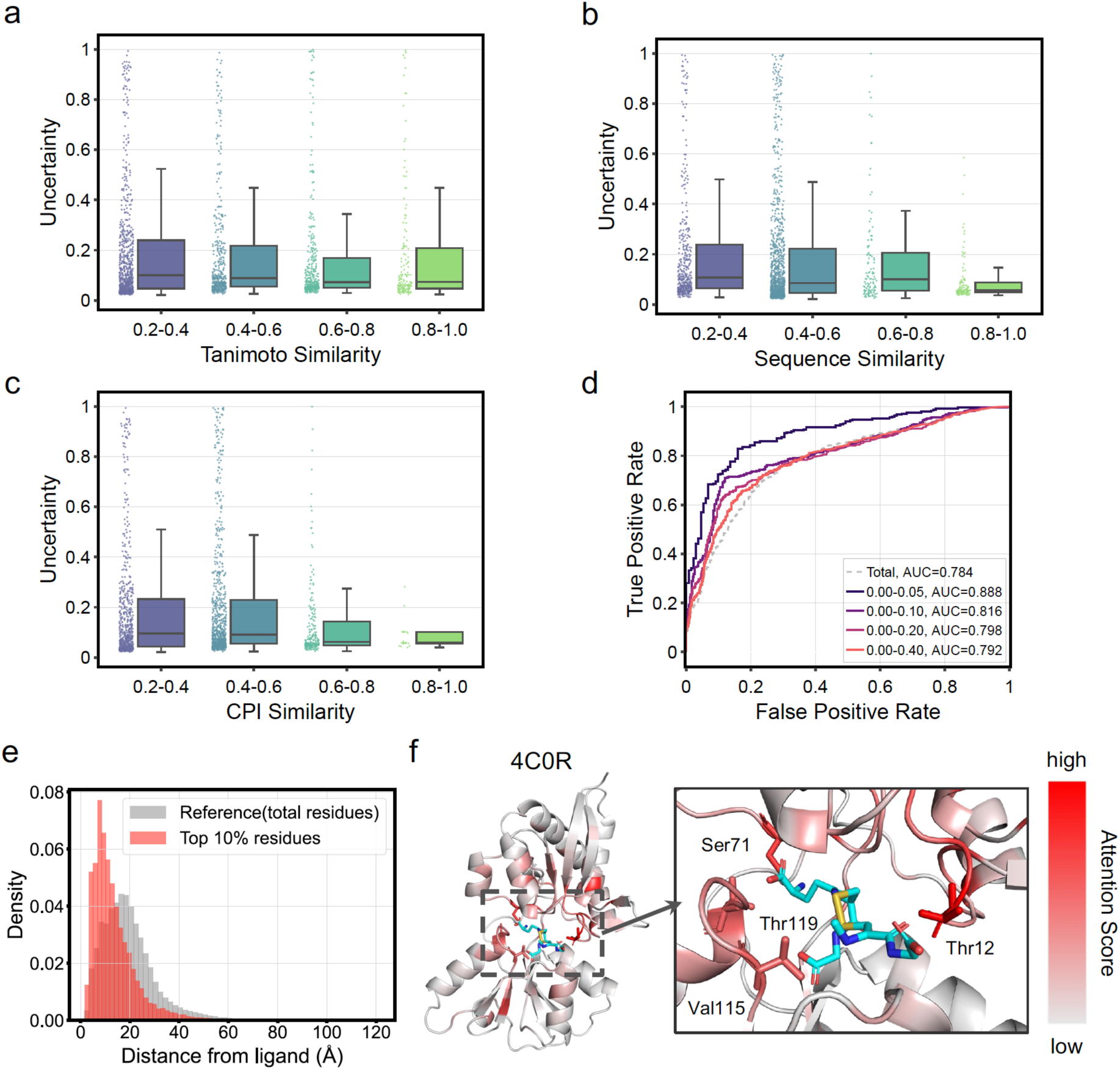
Uncertainty estimation and visualization of attention map to facilitate drug development. **a–c)** Distributions of uncertainty scores based on Tanimoto, sequence, and CPI similarities for the BindingDB dataset. The test set was grouped based on the similarity to the training set in increments of 0.2, and the distribution for each group was represented with a Boxplot. Plots for intervals with no corresponding samples were not displayed. **d)** Receiver operating characteristic (ROC) curve stratified based on uncertainty scores. The test set was divided into groups based on uncertainty score ranges of 0.0–0.05, 0.0–0.10, 0.0–0.20, and 0.0–0.40, and ROC curves were plotted for each group. **e)** Distance distributions in residues with high attention. The histogram in red shows the distance distribution of residues with high attention scores (top 10%), while that in grey represents a reference distribution for all residues. **f)** Attention visualization for Glutathione disulfide (GSSG) and Glutathione transporter (GshT) (Protein Data Bank (PDB) entry: 4C0R). The left panel represents the tertiary complex structure. The right panel is an enlarged view of the regions indicated by the dashed rectangle in the middle panel. The atoms and residues with high attention scores are highlighted in red. In the tertiary structure, GSSG and GshT are represented by ribbon and stick models. The key residues forming interactions with GSSG in the experimental structure, Thr12, Val115, and Thr119, are represented by stick models. Max-pooled attention scores, which were min-max normalized along the sequence direction, were assigned as B-factors. A colour scale ranging from gray90 to red was applied based on these B-factors. The colour scale was set with a minimum value of 0.2 and a maximum value of 1.0.

### ChemGLaM offers a visual understanding of the mechanism of CPIs

The advantage of ChemGLaM also lies in its ability to investigate the relationships between simplified molecular input line entry system (SMILES) compounds and protein sequences through an attention mechanism. This suggests that the visualization of attention weights may offer insights into the molecular mechanisms underlying CPIs. In ChemGLaM, attention matrices are normalised with softmax along the key (residue) direction, enabling the comparison of the attention weights among residues.

Using the model fine-tuned with the PDBbind dataset, the attention weights were mapped onto the tertiary structures. ChemGLaM learns CPIs solely from SMILES compounds and protein sequences. However, residues with high attention weights were primarily distributed around the ligand-binding sites, which were spatially adjacent but sequentially discontinuous (Figure 3e). Despite a certain degree of ambiguity, this observation suggests that the fine-tuned model captures features related to protein folding for ligand recognition. As a representative sample, the glutathione transporter (GshT)-glutathione disulfide (GSSG) complex demonstrated that the highlighted residues agreed well with the interaction pattern observed in the experimentally determined structure (Protein Data Bank (PDB) entry: 4C0R) [36] (Figure 3f). Notably, residues Ser71, Thr12, and Thr119 in GshT are crucial in complex formation by forming hydrogen bonds in the experimental structure. In addition, several residues in the hydrophobic pocket, including Val115, were highlighted in the visualization. Similar results were obtained for other CPIs, where the residues constituting the binding pockets and those surrounding them are highlighted (Supplementary Figure 3), emphasising the effectiveness of providing insights into interaction mechanisms.

### Multimodal framework and large-scale pre-training boost CPI prediction performance

To verify the effectiveness of the multimodal model combining independently pre-trained language models, we conducted ablation studies (Table 1). First, we compared the performance of ChemGLaM without pretraining MoLFormer with that of the original ChemGLaM. Among the four benchmark sets (BindingDB, Davis, PDBbind, and Metz), the performance decreased significantly for BindingDB and Metz compared with that of the original model. This result demonstrates that large-scale pretraining through self-supervised learning is crucial for achieving high generalization. Furthermore, prediction models using only MoLFormer or ESM-2 performed worse than did ChemGLaM, confirming that two independently pre-trained language models synergistically enhance the prediction performance.

We further investigated the relationship between the size of the pre-trained language models and their ability to predict CPIs. We evaluated the CPI prediction performance of ChemGLaM using PLMs with different parameter sizes, ESM-2 (150M, 650M, and 3B parameters) and Prot-T5-XL-UR50 (3B parameters), on the PDBbind dataset. A comparison of the performances based on the RMSE and Pearson correlation revealed that the models using PLMs with larger parameters tended to yield more accurate CPI predictions than did other models, and the use of ESM-2 (3B) led to the best performance among the other PLMs (Extended Data Table 3). This behaviour in the multimodal task is consistent with those in the performances of PLMs alone, drawn upon the metrics, validation perplexity, and scores from the study by Lin et al.^21^ in key benchmarks (LR P@L, CASP14^24^, and CAMEO^25^). These observations indicate that an increase in the parameter size of the pre-trained language models within ChemGLaM would improve CPI prediction performance.

### Large-scale prediction with ChemGLaM provides an enhanced understanding of CPIs

Elucidating the universal relationship between drugs and human proteins is significant for drug development, including drug repurposing, and avoiding unexpected adverse effects. The low computational cost of CPI prediction using ChemGLaM enables large-scale predictions. We conducted comprehensive predictions for all the pairs of 11,455 drugs and 20,434 human proteins, resulting in 234,071,470. For this analysis, we used the ChemGLaM model fine-tuned using 1,392,072 CPI data entries derived from ChEMBL^14^ version 34. Hierarchical clustering of the resulting prediction score profiles depicted a characteristic pattern in the heatmap (Figure 4a), and their clusters demonstrated a strong correlation with the existing annotation. For example, drug Cluster 5 was enriched with drugs categorised under Anatomical Therapeutic Chemical (ATC) code A (alimentary tract and metabolism), whereas drug Cluster 9 predominantly comprised drugs classified under ATC code D (dermatological) (Figure 4b). Similarly, Ser/Thr protein kinases were highly enriched in Cluster 10 (Figure 4c). The score profile within target Cluster 10 revealed a high specificity for particular drugs, possibly reflecting the selectivity of kinase inhibitors. These results indicate that the score profile represents the categorical properties of drugs and targets, providing clues for elucidating the intricate relationships between drugs and proteins. We created comprehensive score profiles, along with uncertainty scores, which are publicly available in a database (https://chemglam.med.kyoto-u.ac.jp/). We believe that this resource enhances the efficiency of drug development by facilitating the discovery of drugs with minimal off-target effects.

**Figure 4:**
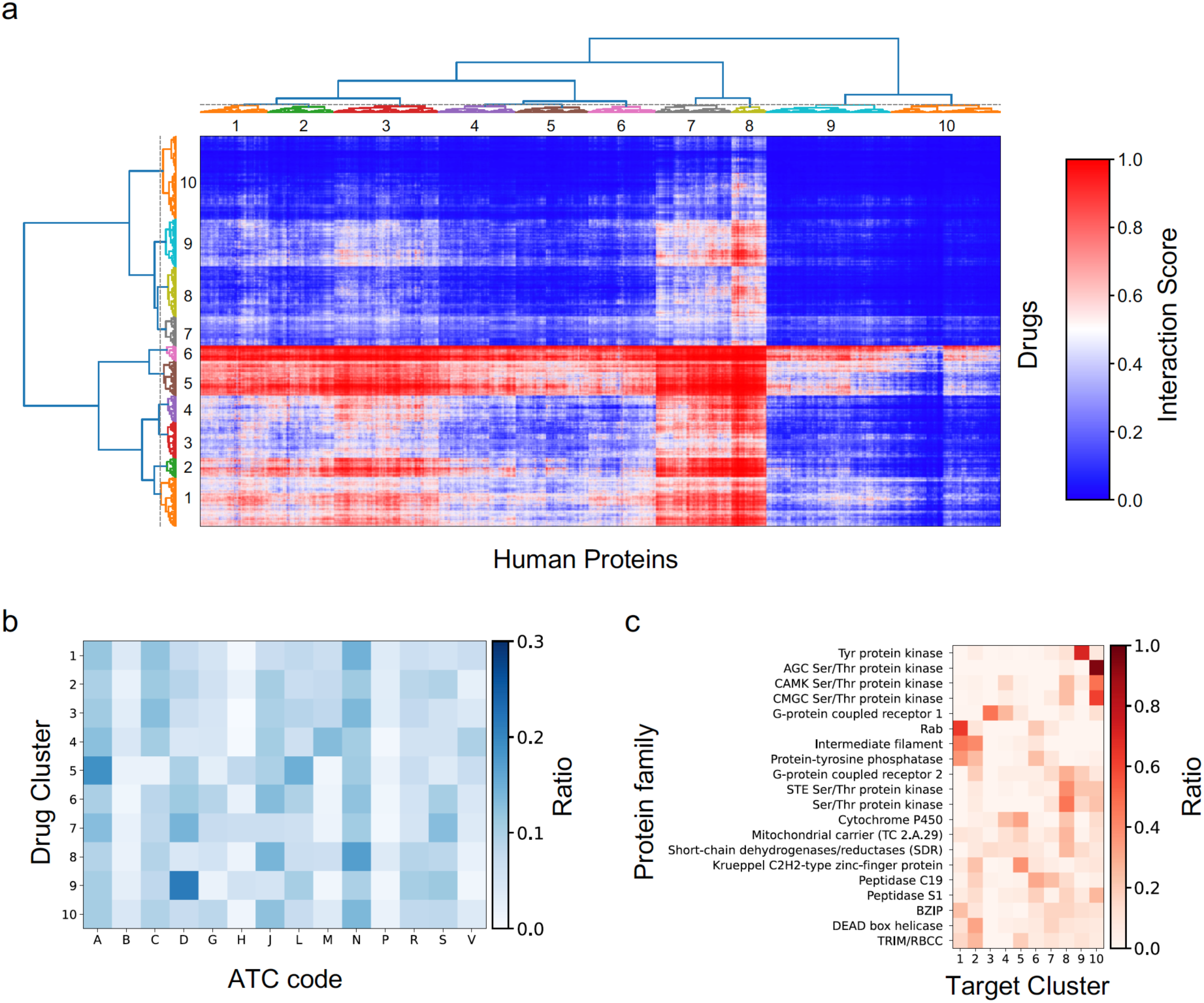
Drug-target interaction (DTI) score profile and clustering analysis. **a)** The interaction scores for all pairs between 2,490 approved drugs and 20,434 human proteins are shown as a heatmap, with the horizontal and vertical axes representing targets and drugs, respectively. Drugs and targets were independently clustered using the hierarchical clustering with Ward’s method. **b)** Enrichment analysis for the drug clusters. The heatmap in the lower left shows the enrichment analysis of the Anatomical Therapeutic Chemical (ATC) classification for the drug clusters. The proportion of ATC classifications within each cluster is colour-coded in blue. **c)** Enrichment analysis for the target clusters. The heatmap in the lower right shows the results of enrichment analysis for the protein families within the target clusters. The proportion of the most abundant 20 protein families belonging to each cluster is colour-coded in red.

### ChemGLaM is applicable to real-world drug screening system

To validate the effectiveness of ChemGLaM in addressing therapeutic challenges, we evaluated its utility in a drug-screening system using induced pluripotent stem cells derived from patients with amyotrophic lateral sclerosis (ALS). In our previous study, we identified compounds capable of rescuing ALS motor neurons from 1,416 compounds including existing drugs for treating various diseases that are commercially available or undergoing clinical testing based on the Motor Neuron Survival Index (MNI)^26^. Among this set, we evaluated the 1,395 compounds that remained after preprocessing using ChemGLaM (see Methods). Separately, to obtain additional data points, we further analysed 212 compounds that were not in the developmental stage at that time. As a previous study reported that the Src/c-Abl signaling pathway is a potential therapeutic target, we conducted comprehensive predictions using ChemGLaM on Src/c-Abl pathway-related proteins (17 proteins: Supplementary Table 2) across a combined set of 1,607 compounds. Subsequently, we performed an enrichment analysis using the maximum cross-protein outputs as the final prediction scores. Good enrichment (AUC = 0.664) was confirmed among the top-scoring compounds (Figure 5a), suggesting that ChemGLaM is effective in exploring candidates in situations where the target molecules are ambiguous. Among the five compounds exhibiting the highest MNIs in the additional data, ChemGLaM successfully identified JAK Inhibitor I and Aurora Kinase Inhibitor II with high prediction scores of 0.982 and 0.990 and low uncertainty scores of 0.035 and 0.019, respectively. The other compounds could not be selected because of their high uncertainty scores (Extended Data Table 4, Figure 5b). For the newly identified hit compounds and their target proteins, selected based on their prediction scores, we predicted their complex structures using HelixFold3^27^, an open-source reproduction of AlphaFold3^28^, and applied attention visualization to the predicted structures. For both candidates, residues spatially adjacent to the predicted binding sites were highlighted (Figure 5c and d). In the predicted structure (Figure 5d) of Aurora Kinase Inhibitor II and Protein kinase C eta type (KPCL_HUMAN), visualization highlighted residues forming the binding pocket, such as L108, D483, and S365. To further explore the potential target proteins in this assay, we comprehensively evaluated the enrichment of 66 reported targets in ALS treatment^29^ (Supplementary Figure 4, Supplementary Table 3) and found additional proteins such as CSF1R and KIT (Figure 5d) as possible targets for ALS. The candidate compounds may confer neuroprotection by interacting with these proteins. Current attempts suggest the potential of ChemGLaM for exploring promising compounds, target proteins, and their interaction sites, even in cases such as cell-based assays where specific targets are unknown.

**Figure 5:**
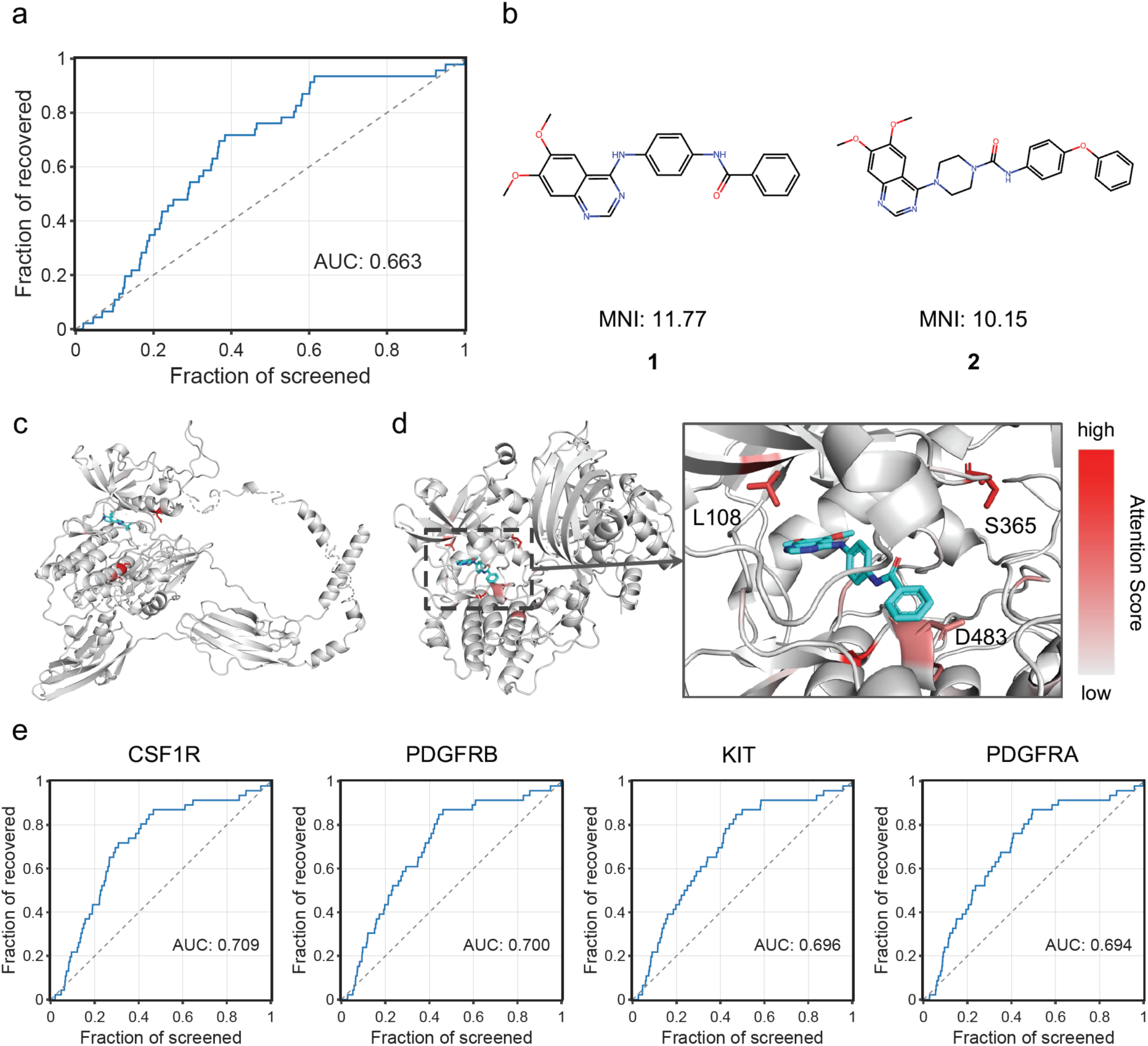
ALS drug screening using ChemGLaM. **a)** Enrichment analysis for drug screening data targeting iPS cells from patients with SOD1-ALS. The enrichment curve was plotted against the maximum scores of cross-protein output scores. **b)** Hit compounds predicted with high scores by ChemGLaM among the newly identified hit compounds from the newly validated library. 1: Aurora Kinase Inhibitor II and 2: PDGFR Tyrosine Kinase Inhibitor III. **c)** Attention visualization for the Drug 1 and its predicted target, protein kinase C eta type. The complex structure is predicted by HelixFold3. The colour scale was set with a minimum value of 0.2 and a maximum value of 1.0. **d)** Attention visualization for the Drug 2 and its predicted target, FGFR2. The complex structure was predicted by HelixFold3. The colour scale was set with a minimum value of 0.2 and a maximum value of 1.0. **e)** The most enriched four target candidates identified by ChemGLaM.

## Discussion

We proposed ChemGLaM, a chemical genomics language model for predicting CPIs. Extensive performance evaluations across multiple CPI datasets demonstrated that our method significantly outperformed other DL-based approaches in predicting unknown CPIs. The ablation study highlighted that this success could be attributed to the integration of multiple language models individually pre-trained on large-scale molecular and protein datasets. This observation provides significant insight that combining such pre-trained large language models is effective for multimodal tasks, such as CPI prediction. The superior performance depends on the scale of the language models used, suggesting that the predictive capability of ChemGLaM can be enhanced using larger models, such as the largest MoLFormer-XL, which is not publicly available presently, or proposed future models.

The visualized cross-attention maps in our model agreed well with the experimentally determined tertiary structures. These results indicate that ChemGLaM learns protein folding and binding events in the 3D space solely from SMILES compound and protein sequences. Ross et al.^20^ discussed how MoLformer learns the spatial relationships between SMILES tokens and demonstrated that the attention map obtained from MoLformer correlated with the contact map derived from the 3D structure of the compound. Similarly, Lin et al.^21^ demonstrated that evolutionarily related sequences in the pre-training dataset and model scaling contribute to improvements in the understanding of protein structure. ChemGLaM may leverage the spatial features acquired by these language models to learn 3D information relevant to CPIs. Elucidating the structures of CPIs is essential for drug development; however, experimental approaches are time-consuming and costly. Visualization of CPI prediction in ChemGLaM could provide structural insight into molecular mechanisms at low computational cost, facilitating the drug discovery process. Notably, although attention visualization tended to highlight the residues adjacent to the bound ligand in the current model, there were many cases where the visualization failed to present the precise experimentally observed interaction patterns. We believe that exploring more sophisticated architectures, including the integration of 3D information and the advancement of foundation models related to compounds and proteins, could enhance the explainability of the model.

Identifying hit compounds as novel targets is essential for drug development, particularly in areas lacking medical needs. ChemGLaM has substantial potential to offer scalable and efficient identification of novel CPIs and their molecular mechanisms. This study could pave the way for a breakthrough in the increasingly complicated process of modern drug development.

## Methods

### ChemGLaM architecture

ChemGLaM comprises three parts: (1) a Compound Encoder, (2) a Protein Encoder, and (3) an Interaction Block. The following section describes the model architecture (Figure 1).

#### Compound encoder: MoLFormer

MoLFormer^20^ is a large-scale chemical language model that learns molecular representation by training on SMILES^30^ sequences of 1.1 billion unlabelled molecules from the PubChem^31^ and ZINC^32^ datasets. MoLFormer-XL, which is the largest MoLFormer variant, outperforms the state-of-the-art chemical foundation models in various prediction tasks. We adopted MoLFormer pre-trained with 10% of the MoLFormer-XL training dataset because the full model is unavailable in the public repository (https://github.com/IBM/molformer).

#### Protein encoder: ESM-2

ESM-2^21^ is a state-of-the-art protein language model for various tasks, such as predicting protein structures and functions. We chose the pre-trained ESM-2-T36-3B-UR50D, which comprised 36 transformer blocks with 3B parameters trained from UniRef50^33^, as a Protein Encoder of the ChemGLaM. Considering computational efficiency, we set the maximum token length (*L*_*p*_) to 2048 to tokenise the amino acid sequences.

#### Interaction block

The interaction Block learns compound-protein interactions using the outputs of the above-mentioned encoders. More concretely, Interaction Block uses the compound embedding *H*_*c*_ ∈ ℝ^*L_c_*×*D_c_*^ and the protein embedding *H*_*p*_ ∈ ℝ^*L_p_*×*D_p_*^ matrices as inputs, where *L*_*c*_ and *L*_*p*_ are token lengths of a SMILES compound and protein sequence, and *D*_*c*_ = 768 and *D*_*p*_ = 2560 are the dimensions of the embeddings. After these embeddings are obtained, the interaction block updates the compound’s representation using multi-head cross-attention mechanisms:

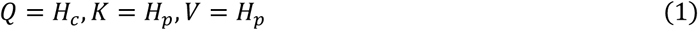

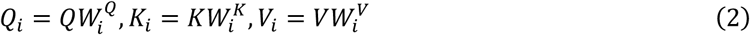

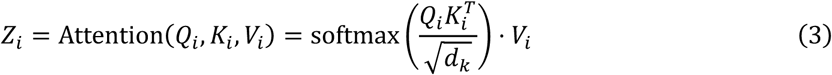

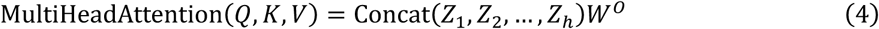

where 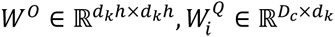, and 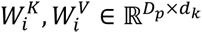 denote weight matrices corresponding to linear layers. We set the number of heads to ℎ = 8, and the number of dimensions *d*_*k*_ for each head was 768/8 = 96. Cross-attention enables the model to focus dynamically on different parts of the two input embedding matrices. This study applied cross attention to compound and protein embeddings to learn what and where to focus on. The cross-attention outputs were added through compound embeddings, followed by layer normalisation^34^.

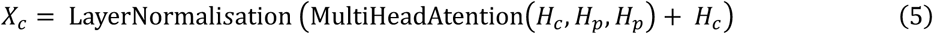

Finally, the average pooled compound features and protein embeddings were concatenated and passed through fully connected layers with skip connections^35^.

### Dataset

Four datasets were prepared to evaluate the effectiveness of ChemGLaM: (1) BindingDB, (2) Davis, (3) PDBbind, and (4) Metz. Supplementary Table 1 presents the statistics for the CPI datasets. We further prepared CPI datasets for predicting all drug pairs from DrugBank and human targets from UniProt by fine-tuning them with a CPI dataset from ChEMBL.

#### BindingDB

BindingDB^15^ is a public, web-accessible database of measured binding affinities that focuses primarily on the interactions between small molecules and proteins. We used a binary classification dataset preprocessed by Gao et al.^36^ with 33,777 positive and 27,493 negative samples.

#### Davis

The Davis^13^ dataset contains data on the binding affinities of several kinase inhibitors to a significant portion of the human kinome. It includes experimentally measured binding constants (Kd values) for kinase-inhibitor pairs. We prepared the same versions of the Davis dataset using DeepDTA^9^. An example was annotated as positive if pKd was more than 5 and negative if pKd was 5 (indicating inactivity in this dataset), resulting in 9,125 and 20,931 positive and negative samples, respectively.

#### PDBbind

PDBbind^16^ is a comprehensive collection of binding affinities of proteins and nucleic acids from the PDB. This database is specially curated for the development of molecular docking algorithms and the study of protein-ligand interactions. A regression dataset was created from the PDBbind version 2020 general set with 18,512 samples.

#### Metz

Metz^6^ is a dataset for CPI research, comprising 1,421 compounds and 156 proteins. Approximately 42% of the compound-protein pairs include binding affinity data as pKi values, representing log-transformed kinase inhibition constants. Consistent experimental conditions and comparable binding affinities of this dataset render it a valuable tool for evaluating predictive models in pharmacological studies. We created a regression dataset from the Metz dataset, which is the same version used by Zhao et al.^6^

### ChEMBL

Activity data targeting single proteins were collected using ChEMBL^14^ version 34. We extracted CPI data with an assay type of ‘B’ and standard units of concentration (uM, nM, or mM) or %. For CPI data with concentration units, we selected entries where the standard types were ‘IC50’, ‘Inhibition’, ‘Activity’, ‘INH’, ‘Ki’, ‘Kd’, or ‘EC50’. For CPI data with % units, we extracted data with standard values ranging from −10 to 110. After merging these datasets, we standardised the SMILES molecules (canonicalisation and largest fragment selection implemented with RDKit) by removing molecules with fewer than four heavy atoms to exclude overly small or inorganic molecules. Additionally, using the method of Ross et al.^20^, we selected only molecules with token sizes less than 200. The resulting dataset was used to train ChemGLaM, which was used to predict the interactions between all drugs and human proteins.

### All pairs between drugs and human proteins

The drugs were extracted from DrugBank version 5.1.12^37^. Human proteins were extracted using UniProt^38^. We created a dataset for all the pairs of drugs and targets. SMILES molecules from DrugBank were preprocessed similarly to those of ChEMBL. The ATC classification of these drugs was extracted from the KEGG database^39^.

### Training and evaluation

#### Training details

Training ChemGLaM involves fine-tuning the Compound Encoder, which is the pre-trained MoLFormer-XL, along with the interaction blocks. We kept the Protein Encoder fixed, considering that ESM-2, which has 3B parameters, was excessively large for the CPI datasets. ChemGLaM was trained using the Adam^40^ optimiser at learning rate = 5 × 10^−6^ over 10 epochs. The batch size was set to 16 for all datasets. This training was performed using a single NVIDIA A100 40GB.

For the loss function, we used cross-entropy loss for the classification tasks, whereas the mean squared error (MSE) loss was used for the regression tasks. When EDL was required for classification tasks, we applied the EDL loss, as subsequently described in Equation (12).

#### Model evaluation: Zero-shot prediction

To verify the generalization performance for unknown CPIs, we conducted model evaluations akin to zero-shot predictions unless otherwise stated. Moreover, we implemented 5-fold cross-validation on test data where both compounds and proteins were unknown, preventing the overestimation of model performance commonly observed in random splits. A diagram of the split datasets with CPI matrices is displayed in Supplementary Figure 1.

#### Baseline models

Hereafter, baseline models are used for model evaluation. To evaluate the effectiveness of self-supervised pretraining in CPI prediction, we primarily focused on chemical genomics-based approaches as baseline models: GraphDTA^5^, TransformerCPI^4^, HyperAttentionDTI^41^, PerceiverCPI^42^, AttentionDTA^6^, Kang et al.^43^, and Conplex^44^. For the PDBbind dataset, we also compared it with a structure-based scoring function such as Vina^11^. We used the Vina scoring function to determine the affinity of the crystal structures from the PDBbind dataset. For models implemented only for a classification or regression task, we implemented training scripts with binary cross-entropy and MSE as the loss functions for the classification and regression tasks, respectively.

### Uncertainty estimation using EDL

EDL is a framework that incorporates uncertainty estimation into the prediction process of deep learning models^22^. Instead of outputting the point estimates of class probabilities, EDL models output the parameters of a Dirichlet distribution as prior distributions. This approach enables a more comprehensive representation of the uncertainty in predictions.

Assuming a K-class classification challenge, the probability density function for the Dirichlet distribution is formulated as follows:

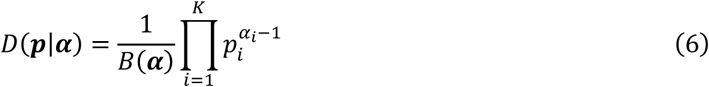

where ***α*** = [*α*_1_,…, *α*_*K*_], (*α*_*i*_ > 1 for all *i* = 1,…, *K*) is a parameter for the Dirichlet distribution, and *B*(***α***) denotes a multinominal beta function.

In EDL, instead of producing regular predictive probabilities, the model outputs the parameters α of the Dirichlet distribution. This distribution represents the uncertainty of the class probabilities. The expected probability of belonging to the kth class is computed as the expected value of the corresponding Dirichlet distribution.

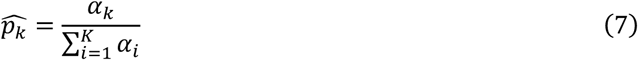

In a binary classification case (*K* = 2) where we predicted whether a compound-protein pair exhibits activity, the probability of activity (second element) was given by the expected probability:

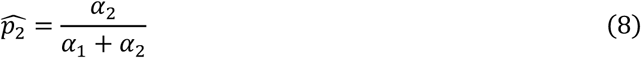

Consequently, the uncertainty *u* was calculated as follows:

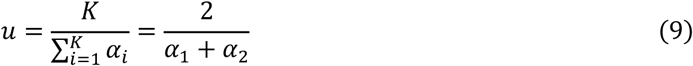

From several loss functions proposed by Sensoy et al. (2018)^22^, we selected the Bayes risk with cross-entropy loss related to the *i*-th sample:

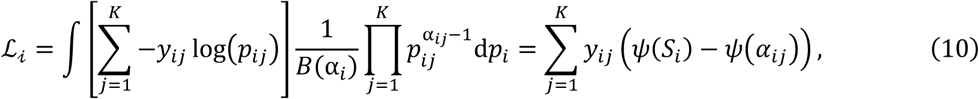

where *ψ*(x) is the digamma function, and 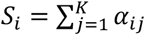. Although they selected Bayes risk with squared error loss, we chose cross-entropy loss because it generated excessively robust evidence in our experiments. To reduce the total evidence to zero if the sample cannot be correctly classified, the EDL loss incorporates a Kullback–Leibler (KL) divergence term:

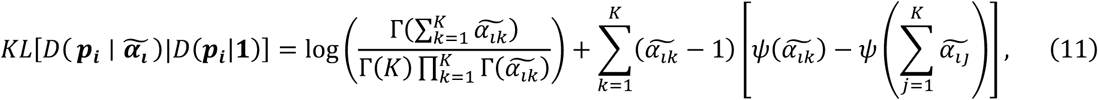

where **1** is the parameter vector of *K* ones, Γ(x) is the gamma function, and 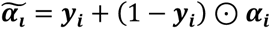 are the Dirichlet parameters after the removal of the non-misleading evidence from a predicted parameter ***α***_***i***_. The final loss function with this regularising term for *N* samples is:

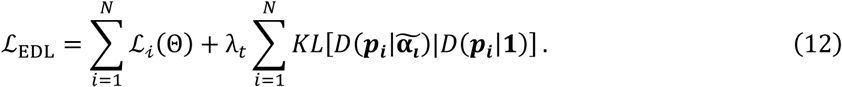

In contrast to the original study^22^, we set the annealing coefficient *λ*_*t*_ as 1 through all the epochs for stable learning.

### Calculating similarities

To evaluate the model performance and uncertainties based on the similarities between the test and training sets, we calculated three types of similarities: Tanimoto, sequence (local alignment score), and their harmonic mean for compound, protein, and CPI similarities. The Tanimoto similarity was calculated using ECFP4^45^ with 2048 bits. Sequence similarity was calculated using a normalised version of the Smith–Waterman^46^ score, where the sequence similarity between two proteins *g* and *g*′ is given by:

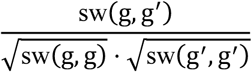

where sw(.,.) represents the original Smith–Waterman score. To divide the test sets based on these similarities, we calculated the pairwise similarities between the training and test sets and assigned the maximum score to each test set.

### Attention visualization

The ChemGLaM models were fine-tuned for the PDBbind dataset by randomly splitting the training and test sets. Attention maps (Equation (3)) for compound-protein pairs were extracted from the fine-tuned models. These maps contain attention scores with dimensions *L*_*c*_ × *L*_*p*_ × ℎ, where *L*_*c*_ and *L*_*p*_ are the token lengths of the compounds and proteins, respectively, and ℎ = 8 is the number of heads. These maps were first averaged through the head dimensions, resulting in attention maps of size *L*_*c*_ × *L*_*p*_. The max-pooled scores for these attention maps were subsequently mapped onto protein residues.

### Ethics

The use of human induced pluripotent stem cells (iPSCs) was approved by the Ethics Committees of Kyoto University. All experiments were performed in accordance with approved guidelines.

### Drug screening using ALS motor neurons and evaluation with ChemGLaM

We conducted high-throughput compound screening using ALS patient iPSC derived motor neurons^26^. The libraries used in the compound screens were MicroSource US Drugs (MicroSource Discovery Systems), MicroSource International Drugs (MicroSource Discovery Systems), and kinase inhibitors from EMD and Selleck Chemicals. The data from previously published screening of existing and clinically tested drugs (n=1,416)^26^, of which 1,395 compounds passed SMILES standardization using the same method as ChEMBL, were evaluated using ChemGLaM. Additional 212 compounds that were not in the developmental stage at the time were screened using the same experimental protocol as the previous study^26^. These compounds were evaluated experimentally, and the results were compared with the ChemGLaM predictions.

## Data availability

The benchmark sets for model evaluation (BindingDB, Davis, PDBbind, and Metz) are available on GitHub (https://github.com/clinfo/ChemGLaM). For the ChEMBL dataset, the source code for the data preprocessing is available in the repository. Large-scale prediction results for all drug pairs (from DrugBank) and human proteins (from UniProt) are available at https://chemglam.med.kyoto-u.ac.jp/.

## Code availability

The source code of ChemGLaM is available on GitHub (https://github.com/clinfo/ChemGLaM), together with usage documentation and the setup environment.

## Author contributions

T. Koyama conceived the present idea and contributed to the algorithm implementation. Y.O., S.M., and T. Kato supervised the project. H.T., R.O., and T. Koyama conducted the model evaluation. K.Y. and T. Koyama were responsible for large-scale training and inference. A.H., H. Iwata, S.M. and T. Koyama contributed to the database construction. K.I. and H. Inoue conducted the drug screening experiment. T. Koyama, S.M., R.K., and Y.O. wrote the manuscript. All authors contributed to discussions and proofreading.

## Acknowledgements

This research was supported by JSPS KAKENHI Grant Number JP24KJ1510 to T. Koyama, 22K06112 to S.M., a Grant-in-Aid for Transformative Research Areas (A) “Latent Chemical Space” [JP24H01771] for H.Iwata. from the Ministry of Education, Culture, Sports, Science and Technology, Japan, and the Japan Agency for Medical Research and Development (AMED) under Grant Number JP22bm0804034, JP23bm1423014, and JP23bm1223013 to H. Inoue. The cartoons of the Petri dish and the protein in Figure 1 were obtained from The Togo Picture Gallery (copyright 2016 DBCLS TogoTV / CCBY-4.0). OpenAI ChatGPT was used to improve the wording of some paragraphs, but not to generate new content. We would like to thank Editage (www.editage.jp) for English language editing.

---

**Extended Data Table 1:** Performance evaluation for the classification tasks (BindingDB and Davis)

| Dataset | Method | AUROC (Std) | AUPRC (Std) | Accuracy (Std) | F1 (Std) |
| --- | --- | --- | --- | --- | --- |
| **BindingDB** | GraphDTA | 0.689 (0.063) | 0.715 (0.107) | 0.572 (0.084) | 0.649 (0.073) |
|  | TransformerCPI | 0.681 (0.035) | 0.716 (0.063) | 0.627 (0.052) | 0.588 (0.032) |
|  | HyperAttentionDTI | 0.792 (0.041) | 0.803 (0.055) | 0.695 (0.060) | 0.663 (0.092) |
|  | PerceiverCPI | 0.678 (0.043) | 0.675 (0.116) | 0.613 (0.063) | 0.535 (0.075) |
|  | AttentionDTA | 0.779 (0.014) | 0.788 (0.063) | 0.710 (0.025) | 0.708 (0.033) |
|  | Kang et al. | 0.803 (0.021) | 0.803 (0.072) | 0.733 (0.043) | 0.715 (0.074) |
|  | ConPLex | 0.813 (0.027) | 0.817 (0.031) | 0.739 (0.033) | 0.732 (0.044) |
|  | ChemGLaM | **0.829 (0.042)** | **0.827 (0.055)** | **0.754 (0.039)** | **0.756 (0.021)** |
| **Davis** | GraphDTA | 0.661 (0.072) | 0.470 (0.131) | 0.694 (0.049) | 0.382 (0.150) |
|  | TransformerCPI | 0.664 (0.072) | 0.460 (0.165) | 0.705 (0.104) | 0.426 (0.174) |
|  | HyperAttentionDTI | 0.671 (0.045) | 0.499 (0.098) | 0.614 (0.067) | 0.486 (0.084) |
|  | PerceiverCPI | 0.671 (0.083) | 0.464 (0.124) | 0.707 (0.032) | 0.410 (0.158) |
|  | AttentionDTA | 0.679 (0.039) | 0.511 (0.076) | 0.596 (0.072) | 0.484 (0.075) |
|  | Kang et al. | 0.704 (0.038) | 0.499 (0.053) | 0.661 (0.031) | 0.502 (0.070) |
|  | ConPLex | 0.666 (0.068) | 0.304 (0.084) | **0.791 (0.017)** | 0.295 (0.105) |
|  | ChemGLaM | **0.728 (0.047)** | **0.561 (0.119)** | 0.738 (0.022) | **0.493 (0.137)** |

AUROC: Area under the receiver operating characteristic curve.
AUPRC: Area under the precision-recall curve.
Std: Standard deviation.

**Extended Data Table 2:** Performance evaluation for the regression tasks (PDBbind and Metz dataset)

| Dataset | Method | RMSE (Std) | Pearson (Std) | Spearman (Std) | MAE (Std) |
| --- | --- | --- | --- | --- | --- |
| **PDBbind** | Vina | - | - | 0.511 (0.037) | - |
|  | GraphDTA | 1.755 (0.137) | 0.441 (0.059) | 0.449 (0.037) | 1.386 (0.095) |
|  | TransformerCPI | 1.733 (0.050) | 0.474 (0.032) | 0.468 (0.024) | 1.369 (0.041) |
|  | HyperAttentionDTI | 1.580 (0.039) | 0.542 (0.026) | 0.539 (0.021) | 1.237 (0.035) |
|  | PerceiverCPI | 1.687 (0.061) | 0.462 (0.052) | 0.450 (0.045) | 1.337 (0.044) |
|  | AttentionDTA | 1.615 (0.075) | 0.523 (0.047) | 0.509 (0.044) | 1.283 (0.057) |
|  | Kang et al. | 1.743 (0.034) | 0.569 (0.021) | 0.562 (0.027) | 1.374 (0.044) |
|  | ConPLex | 1.726 (0.016) | 0.503 (0.027) | 0.519 (0.023) | 1.368 (0.028) |
|  | ChemGLaM | **1.521 (0.047)** | **0.600 (0.026)** | **0.596 (0.024)** | **1.198 (0.031)** |
| **Metz** | GraphDTA | 1.314 (0.108) | 0.363 (0.054) | 0.330 (0.056) | 1.034 (0.111) |
|  | TransformerCPI | 0.879 (0.045) | 0.442 (0.048) | 0.400 (0.052) | 0.705 (0.035) |
|  | HyperAttentionDTI | 0.907 (0.082) | 0.419 (0.052) | 0.368 (0.046) | 0.693 (0.048) |
|  | PerceiverCPI | 0.913 (0.032) | 0.399 (0.071) | 0.378 (0.071) | 0.704 (0.026) |
|  | AttentionDTA | 0.858 (0.046) | 0.482 (0.023) | 0.419 (0.033) | 0.655 (0.033) |
|  | Kang et al. | 0.993 (0.046) | 0.515 (0.042) | 0.461 (0.037) | 0.831 (0.043) |
|  | ConPLex | 1.553 (0.060) | 0.358 (0.022) | 0.342 (0.035) | 1.195 (0.043) |
|  | ChemGLaM | **0.812 (0.029)** | **0.544 (0.025)** | **0.485 (0.030)** | **0.643 (0.026)** |

RMSE: Root mean squared error
MAE: Mean absolute error
Std: Standard deviation

**Extended Data Table 3:**
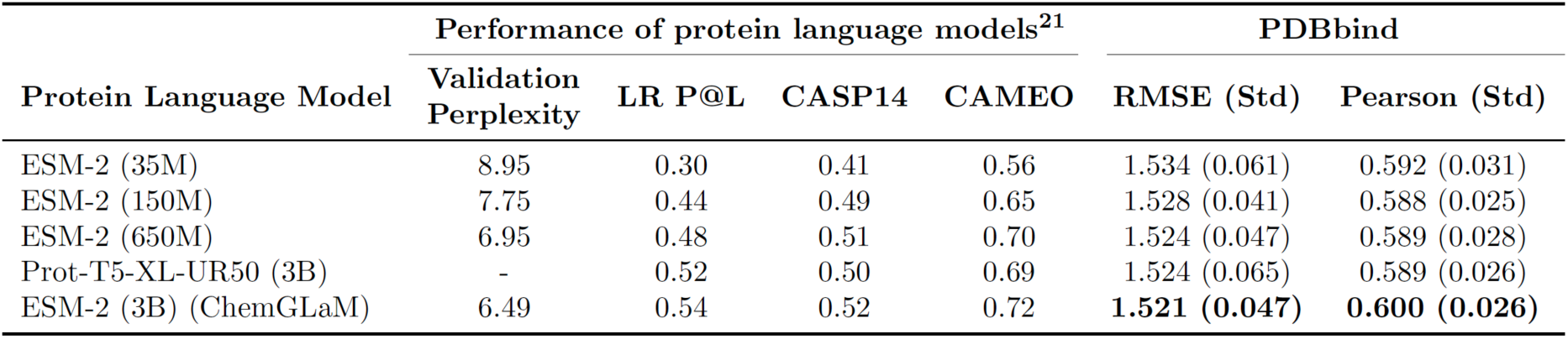
Relation between the performance of protein language models and that of CPI prediction for PDBbind dataset.

| Protein Language Model | Performance of protein language models<sup>21</sup> |  |  |  | PDBbind |  |
| --- | --- | --- | --- | --- | --- | --- |
|  | Validation Perplexity | LR P@L | CASP14 | CAMEO | RMSE (Std) | Pearson (Std) |
| ESM-2 (35M) | 8.95 | 0.30 | 0.41 | 0.56 | 1.534 (0.061) | 0.592 (0.031) |
| ESM-2 (150M) | 7.75 | 0.44 | 0.49 | 0.65 | 1.528 (0.041) | 0.588 (0.025) |
| ESM-2 (650M) | 6.95 | 0.48 | 0.51 | 0.70 | 1.524 (0.047) | 0.589 (0.028) |
| Prot-T5-XL-UR50 (3B) | - | 0.52 | 0.50 | 0.69 | 1.524 (0.065) | 0.589 (0.026) |
| ESM-2 (3B) (ChemGLaM) | 6.49 | 0.54 | 0.52 | 0.72 | **1.521 (0.047)** | **0.600 (0.026)** |

**Extended Data Table 4:**
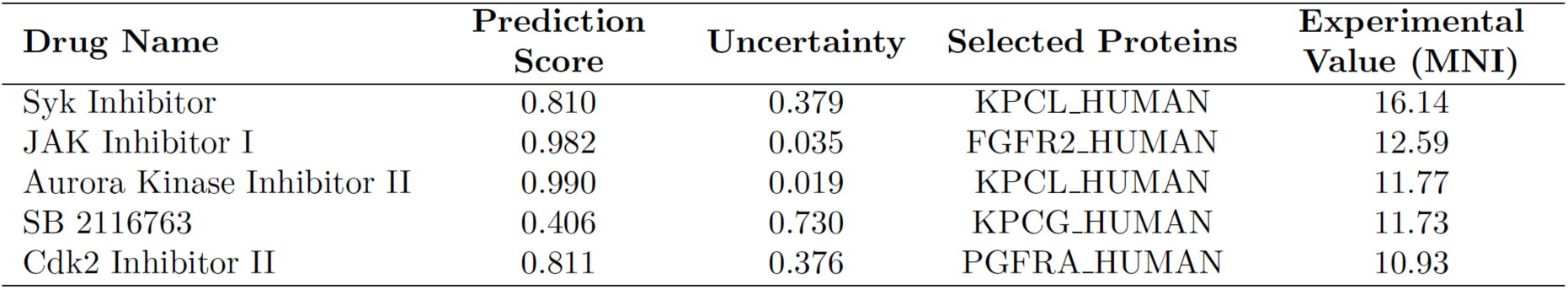
Predicted and experimental values of the five most active compounds among the newly obtained hit compounds.

## Supplementary Information

**Supplementary Table 1:** Summary of CPI datasets.

| Name | Task | Compounds | Proteins | Interactions | Positives | Negatives |
| --- | --- | --- | --- | --- | --- | --- |
| BindingDB | Classification | 49,752 | 813 | 61,270 | 33,777 | 27,493 |
| Davis | Classification | 68 | 442 | 30,056 | 9,125 | 20,931 |
| PDBbind | Regression | 14,694 | 3,705 | 18,512 | - | - |
| Metz | Regression | 1,423 | 170 | 35,259 | - | - |

**Supplementary Figure 1:**
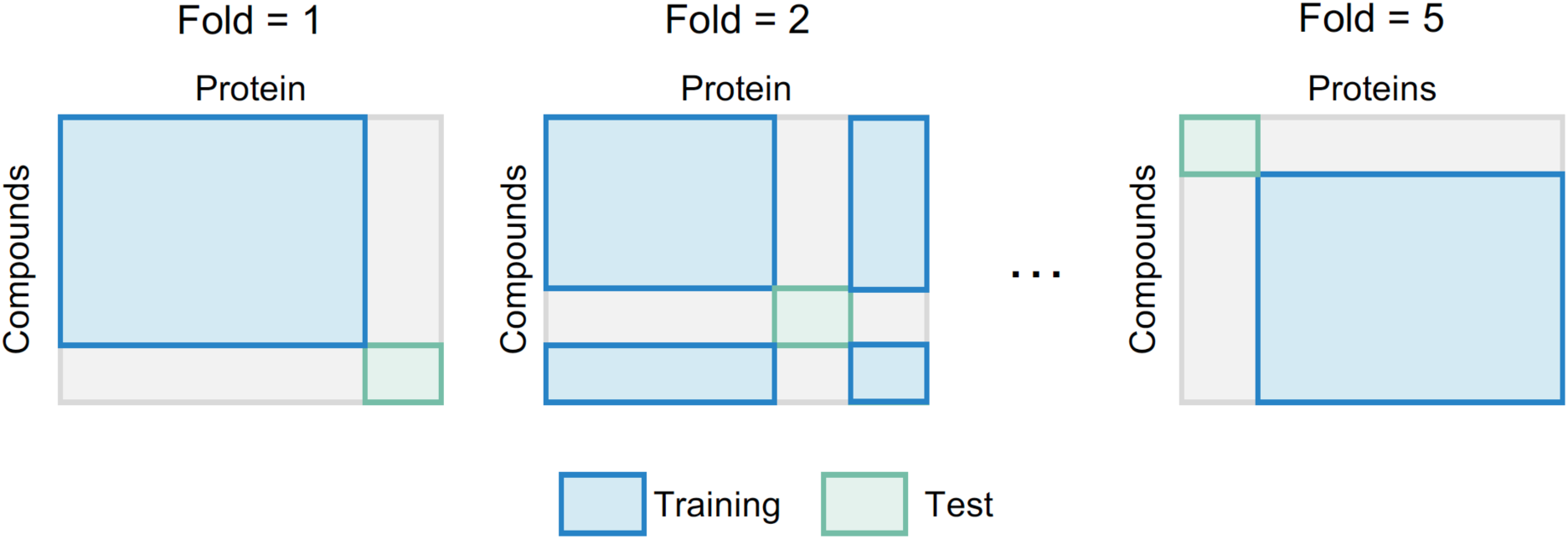
Schematic illustration of zero-shot prediction. The whole dataset was divided into training and test sets so that both datasets don’t share any compounds and proteins. The vertical axis of the matrices represents compounds, and the horizontal axis represents proteins.

**Supplementary Figure 2:**
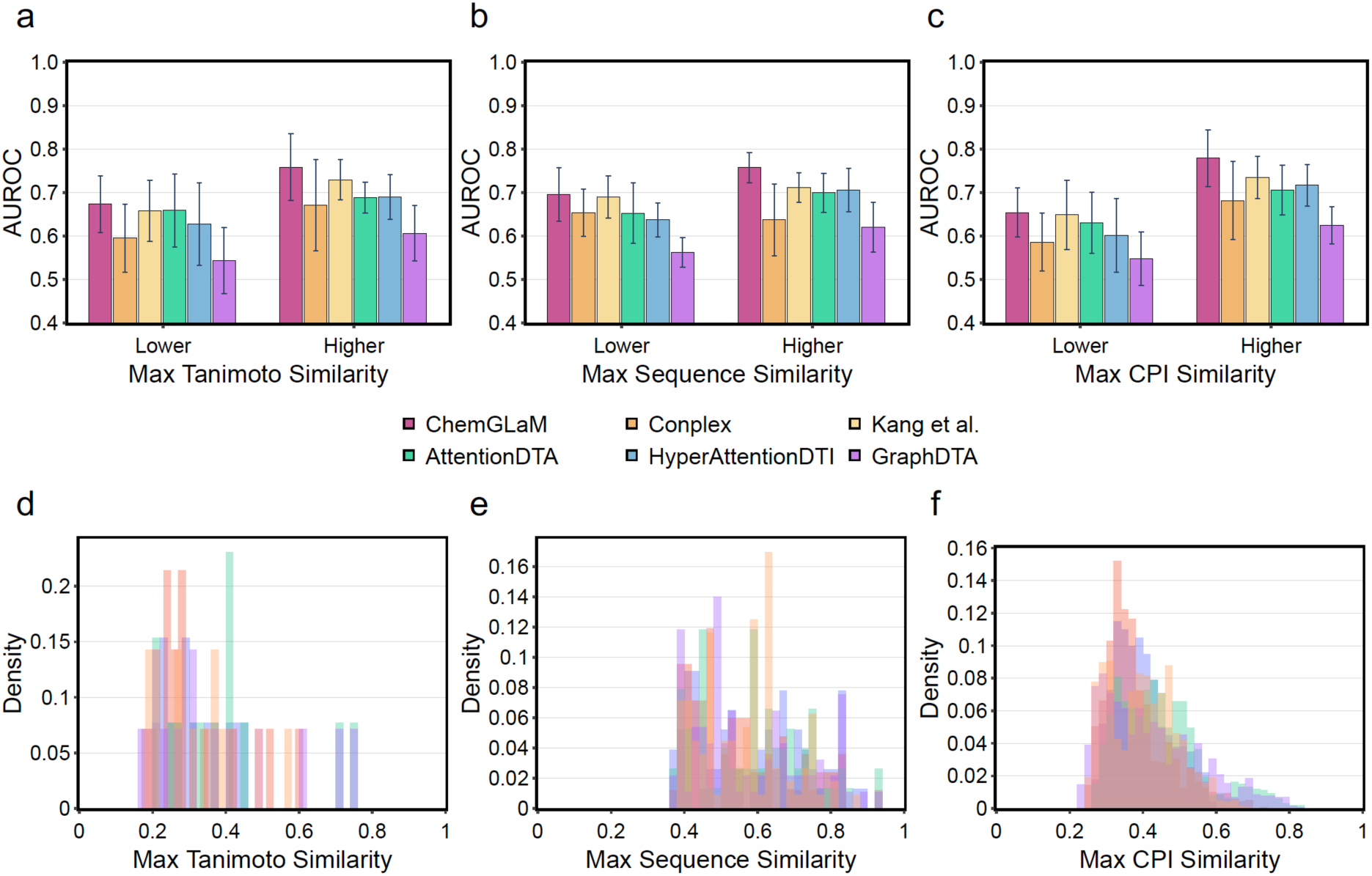
Performance evaluation of ChemGLaM in zero-shot prediction. **a-c)** The average AUROC scores of 5-fold cross-validation for the Davis dataset. The test set in each fold is divided into two groups based on the maximum similarities with the training set: close and distant pairs. The similarities are calculated via three approaches: Tanimoto, sequence, and CPI similarities. The CPI similarity is a harmonic mean of Tanimoto and sequence similarities. The bars on the left side represent the AUROC scores for the distant pairs (lower similarity) and those on the right side represent the scores for the close pairs (higher similarity). **d-f)** Distributions of maximum Tanimoto, sequence, and CPI similarities in five folds are represented. The distributions in each fold are indicated by different colours.

**Supplementary Figure 3:**
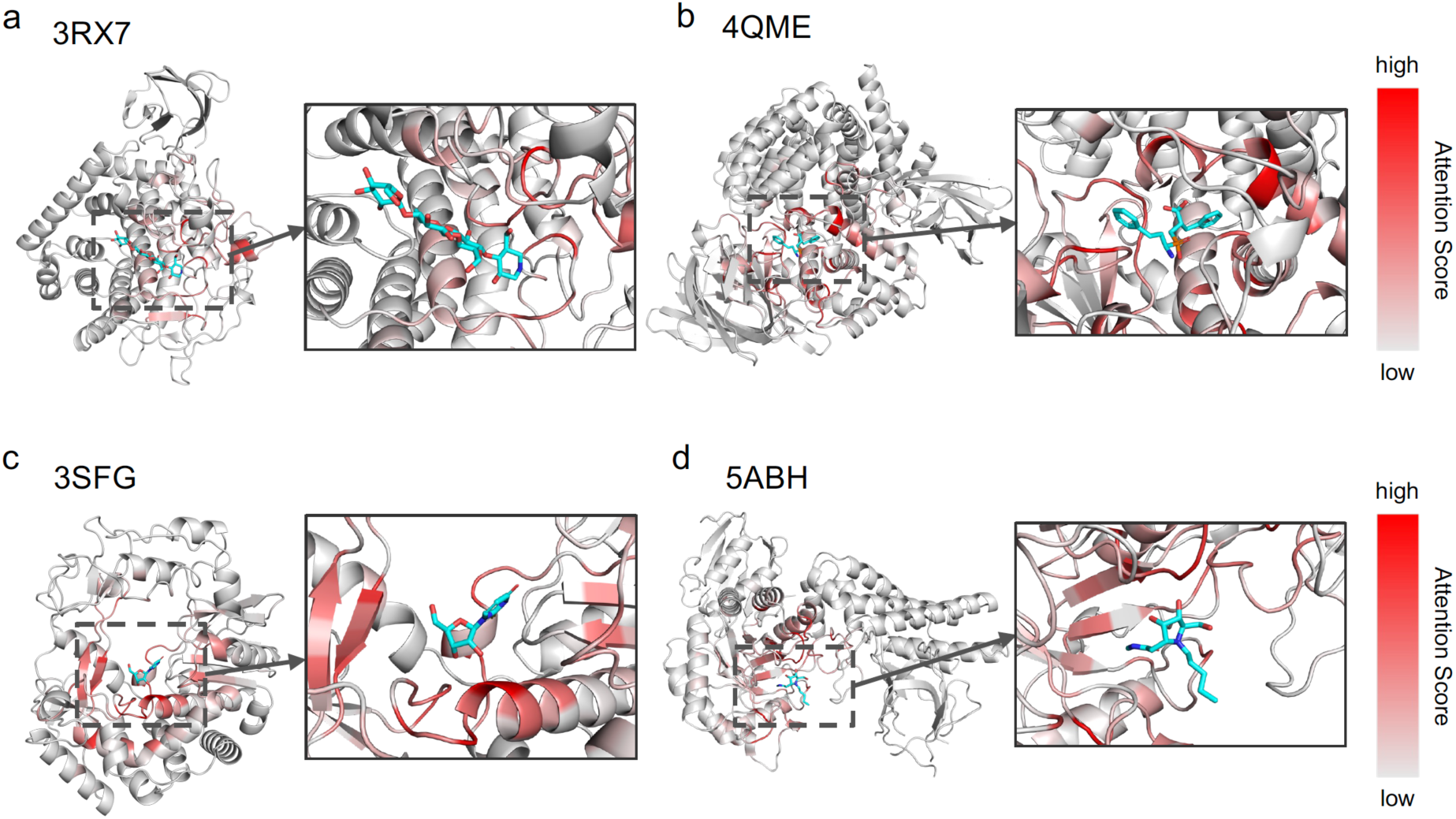
Visualizing attention maps on other complex structures. Attention visualization for **a**) AaCel9A in complex with cellotetraose-like isofagomine (PDB entry = 3RX7), **b**) Aminopeptidase N in complex with the phosphinic dipeptide analogue LL-(R, S)-hPheP[CH2]Phe (PDB entry = 4QME), **c**) murine norovirus RNA dependent RNA polymerase in complex with 2thiouridine(2TU) (3SFG), and **d)** GH84 with ligand (PDB entry = 5ABH). Max-pooled attention scores, which were min-max normalized along the sequence direction, were assigned as B-factors. A color scale ranging from gray90 to red was then applied based on these B-factors. The color scale was set with a minimum value of 0.2 and a maximum value of 1.0.

**Supplementary Table 2:** Protein list used in current study as related to Src/c-Abl pathway.

| UniProt ID | Protein names |
| --- | --- |
| P00519 | Tyrosine-protein kinase ABL1 |
| Q9UM73 | ALK tyrosine kinase receptor |
| P11362 | Fibroblast growth factor receptor 1 |
| P21802 | Fibroblast growth factor receptor 2 |
| P22607 | Fibroblast growth factor receptor 3 |
| P22455 | Fibroblast growth factor receptor 4 |
| P17252 | Protein kinase C alpha type |
| P05771 | Protein kinase C beta type |
| P05129 | Protein kinase C gamma type |
| P24723 | Protein kinase C eta type |
| P08581 | Hepatocyte growth factor receptor |
| P16234 | Platelet-derived growth factor receptor alpha |
| P09619 | Platelet-derived growth factor receptor beta |
| P12931 | Proto-oncogene tyrosine-protein kinase Src |
| P17948 | Vascular endothelial growth factor receptor 1 |
| P35968 | Vascular endothelial growth factor receptor 2 |
| P35916 | Vascular endothelial growth factor receptor 3 |

**Supplementary Figure 4:**
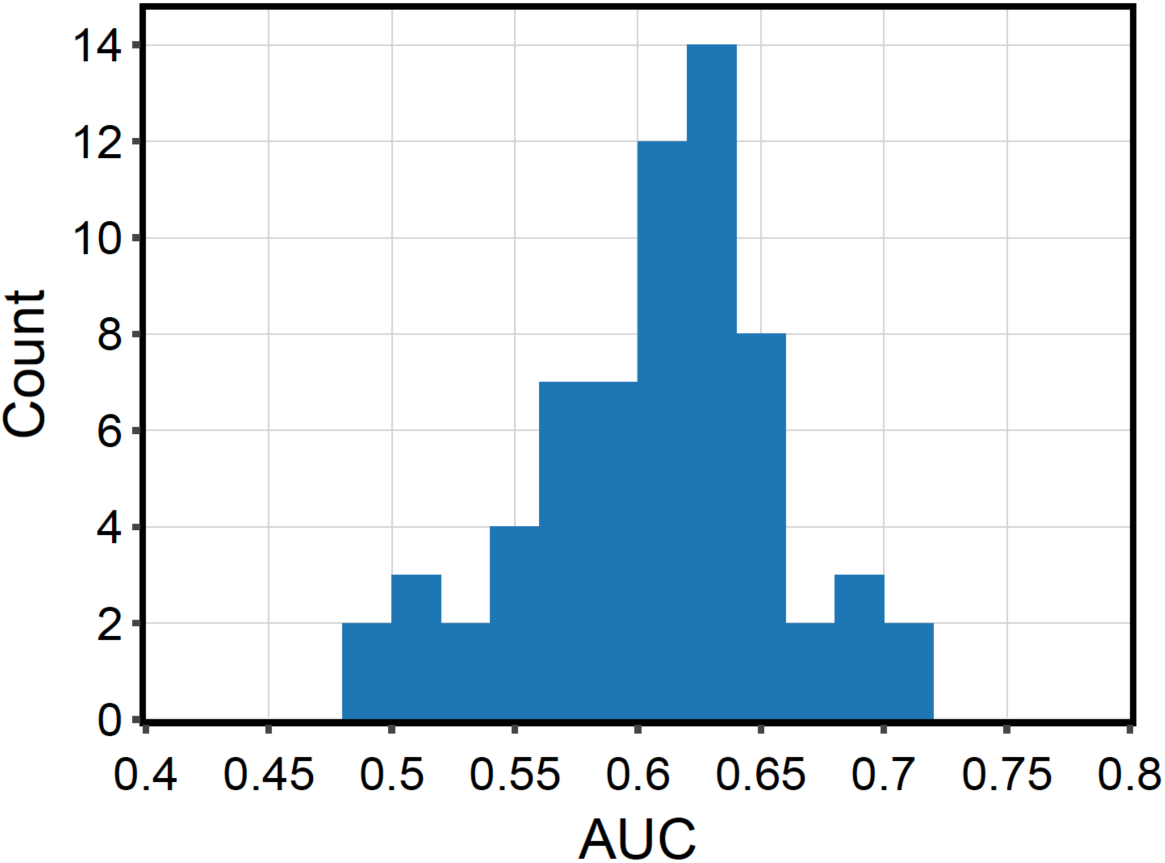
Distribution of AUC values for hit enrichment curves of 66 reported targets in ALS treatment ^2^.

**Supplementary Table 3:**
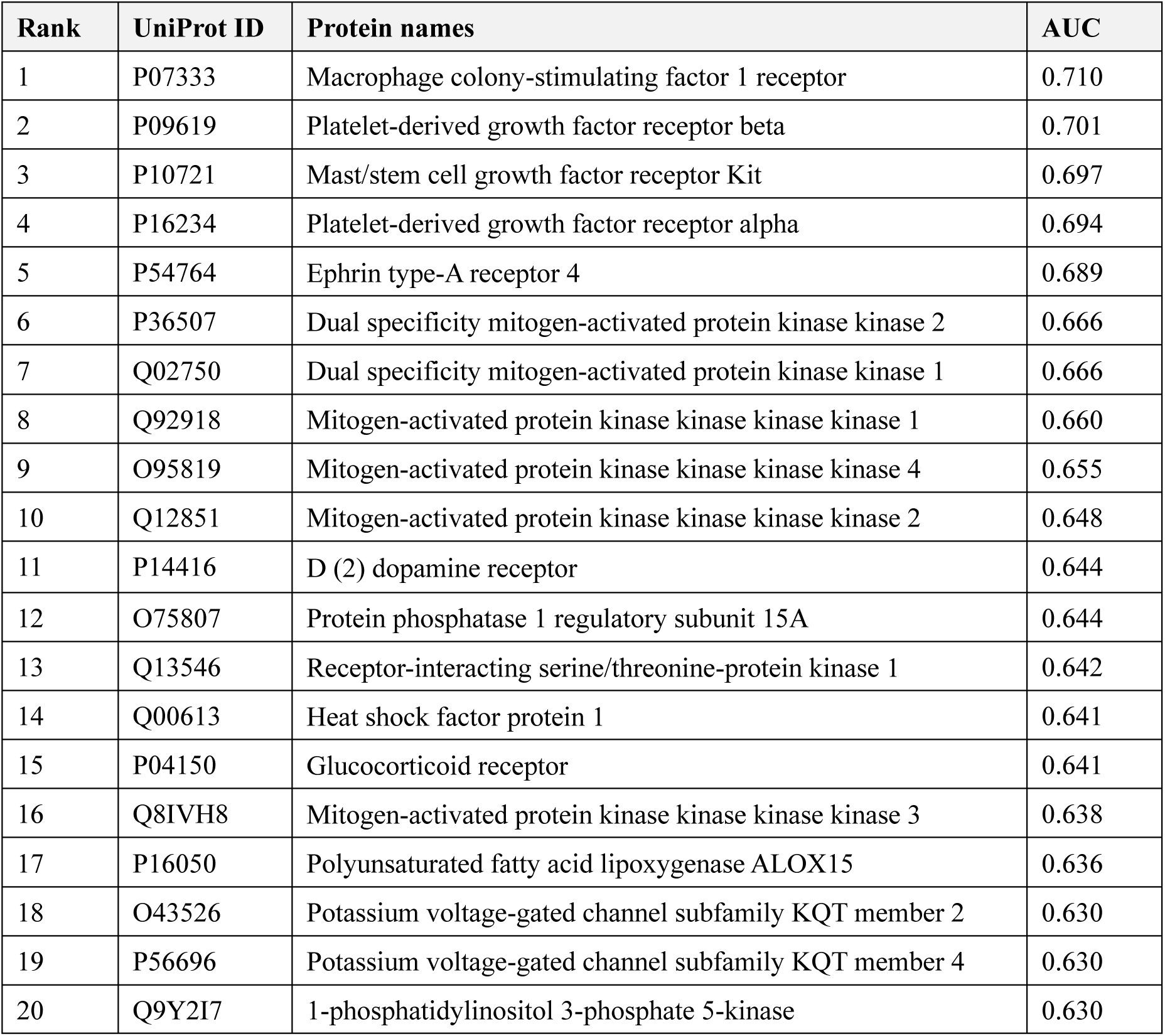
Most enriched 20 target candidates of 66 reported targets in ALS treatment ^1^. The targets are sorted by AUC values for hit enrichment curves.

## References

1. Keiser, M. J. et al. Predicting new molecular targets for known drugs. Nature 462, 175–181 (2009). 10.1038/nature08506, PubMed: 19881490.

2. Drewry, D. H. & Macarron, R. Enhancements of screening collections to address areas of unmet medical need: an industry perspective. Curr. Opin. Chem. Biol. 14, 289–298 (2010). 10.1016/j.cbpa.2010.03.024, PubMed: 20413343.

3. Macarron, R. et al. Impact of high-throughput screening in biomedical research. Nat. Rev. Drug Discov. 10, 188–195 (2011). 10.1038/nrd3368, PubMed: 21358738.

4. Chen, L. et al. TransformerCPI: improving compound–protein interaction prediction by sequence-based deep learning with self-attention mechanism and label reversal experiments. Bioinformatics 36, 4406–4414 (2020). 10.1093/bioinformatics/btaa524, PubMed: 32428219.

5. Nguyen, T. et al. GraphDTA: predicting drug–target binding affinity with graph neural networks. Bioinformatics 37, 1140–1147 (2021). 10.1093/bioinformatics/btaa921, PubMed: 33119053.

6. Zhao, Q. et al. AttentionDTA: drug–target binding affinity prediction by sequence-based deep learning with attention mechanism. IEEE ACM Trans. Comp. Biol. Bioinform. 20, 852–863 (2023). 10.1109/TCBB.2022.3170365, PubMed: 35471889.

7. Kearnes, S., McCloskey, K., Berndl, M., Pande, V. & Riley, P. Molecular graph convolutions: moving beyond fingerprints. J. Comput. Aided Mol. Des. 30, 595–608 (2016). 10.1007/s10822-016-9938-8, PubMed: 27558503.

8. LeCun, Y., Bengio, Y. & Hinton, G. Deep learning. Nature 521, 436–444 (2015). 10.1038/nature14539, PubMed: 26017442.

9. Öztürk, H., Özgür, A. & Ozkirimli, E. DeepDTA: deep drug–target binding affinity prediction. Bioinformatics 34, i821–i829 (2018). 10.1093/bioinformatics/bty593, PubMed: 30423097.

10. Vaswani, A. et al. Attention is all you need. Adv. Neural Inf. Process. Syst. 30 (2017).

11. Trott, O. & Olson, A. J. AutoDock Vina: improving the speed and accuracy of docking with a new scoring function, efficient optimization, and multithreading. J. Comput. Chem. 31, 455–461 (2010). 10.1002/jcc.21334, PubMed: 19499576.

12. Yabuuchi, H. et al. Analysis of multiple compound–protein interactions reveals novel bioactive molecules. Mol. Syst. Biol. 7, 472 (2011). 10.1038/msb.2011.5, PubMed: 21364574.

13. Davis, M. I. et al. Comprehensive analysis of kinase inhibitor selectivity. Nat. Biotechnol. 29, 1046–1051 (2011). 10.1038/nbt.1990, PubMed: 22037378.

14. Gaulton, A. et al. ChEMBL: a large-scale bioactivity database for drug discovery. Nucleic Acids Res. 40, D1100–D1107 (2012). 10.1093/nar/gkr777, PubMed: 21948594.

15. Gilson, M. K., et al. BindingDB in 2015: a public database for medicinal chemistry, computational chemistry and systems pharmacology. Nucleic Acids Res. 44, D1045–D1053 (2016). 10.1093/nar/gkv1072, PubMed: 26481362.

16. Liu, Z. et al. PDB-wide collection of binding data: current status of the PDBbind database. Bioinformatics 31, 405–412 (2015). 10.1093/bioinformatics/btu626, PubMed: 25301850.

17. Krizhevsky, A., Sutskever, I. & Hinton, G. E. ImageNet classification with deep convolutional neural networks. Commun. ACM 60, 84–90 (2017). 10.1145/3065386.

18. Touvron, H. et al. Llama 2: open foundation and fine-tuned chat models. arXiv Preprint ArXiv:2307.09288 (2023). 10.48550/arXiv.2307.09288.

19. Lipinski, C. & Hopkins, A. Navigating chemical space for biology and medicine. Nature 432, 855–861 (2004). 10.1038/nature03193, PubMed: 15602551.

20. Ross, J. et al. Large-scale chemical language representations capture molecular structure and properties. Nat. Mach. Intell. 4, 1256–1264 (2022). 10.1038/s42256-022-00580-7.

21. Lin, Z. et al. Evolutionary-scale prediction of atomic-level protein structure with a language model. Science 379, 1123–1130 (2023). 10.1126/science.ade2574, PubMed: 36927031.

22. Sensoy, M., Kaplan, L. & Kandemir, M. Evidential deep learning to quantify classification uncertainty. Adv. Neural Inf. Process. Syst. 31 (2018).

23. Metz, J. T., et al. Navigating the kinome. Nat. Chem. Biol. 7, 200–202 (2011). 10.1038/nchembio.530, PubMed: 21336281.

24. Kryshtafovych, A., Schwede, T., Topf, M., Fidelis, K. & Moult, J. Critical assessment of methods of protein structure prediction (CASP)-Round XIV. Proteins 89, 1607–1617 (2021). 10.1002/prot.26237, PubMed: 34533838.

25. Haas, J. et al. Continuous Automated Model EvaluatiOn (CAMEO) complementing the critical assessment of structure prediction in CASP12. Proteins 86(Suppl 1), 387–398 (2018). 10.1002/prot.25431, PubMed: 29178137.

26. Imamura, K. et al. The Src/c-Abl pathway is a potential therapeutic target in amyotrophic lateral sclerosis. Sci. Transl. Med. 9, eaaf3962 (2017). 10.1126/scitranslmed.aaf3962, PubMed: 28539470.

27. Liu, L. et al., (2024). Technical report of HelixFold3 for biomolecular structure prediction. arXiv Preprint ArXiv:2408.16975.

28. Abramson, J. et al. Accurate structure prediction of biomolecular interactions with AlphaFold 3. Nature 630, 493–500 (2024). 10.1038/s41586-024-07487-w, PubMed: 38718835.

29. Mead, R. J., Shan, N., Reiser, H. J., Marshall, F. & Shaw, P. J. Amyotrophic lateral sclerosis: a neurodegenerative disorder poised for successful therapeutic translation. Nat. Rev. Drug Discov. 22, 185–212 (2023). 10.1038/s41573-022-00612-2, PubMed: 36543887.

30. Weininger, D. SMILES, a chemical language and information system. 1. Introduction to methodology and encoding rules. J. Chem. Inf. Comput. Sci. 28, 31–36 (1988). 10.1021/ci00057a005.

31. Kim, S., et al. PubChem 2019 update: improved access to chemical data. Nucleic Acids Res. 47, D1102–D1109 (2019). 10.1093/nar/gky1033, PubMed: 30371825.

32. Irwin, J. J. & Shoichet, B. K. Zinc- a free database of commercially available compounds for virtual screening. J. Chem. Inf. Model. 45, 177–182 (2005). 10.1021/ci049714+, PubMed: 15667143.

33. Suzek, B. E. et al. UniRef clusters: a comprehensive and scalable alternative for improving sequence similarity searches. Bioinformatics 31, 926–932 (2015). 10.1093/bioinformatics/btu739, PubMed: 25398609.

34. Ba, J. L., Kiros, J. R. & Hinton, G. E. Layer normalization. arXiv Preprint ArXiv:1607.06450 (2016). 10.48550/arXiv.1607.06450.

35. He, K., Zhang, X., Ren, S. & Sun, J. Proceedings of the IEEE Conference on Computer Vision and Pattern Recognition 770–778.

36. Gao, K. Y. et al. IJCAI 3371-3377.

37. Wishart, D. S. et al. DrugBank 5.0: a major update to the DrugBank database for 2018. Nucleic Acids Res. 46, D1074–D1082 (2018). 10.1093/nar/gkx1037, PubMed: 29126136.

38. UniProt Consortium. UniProt: the universal protein knowledgebase in 2021. Nucleic Acids Res. 49, D480–D489 (2021). 10.1093/nar/gkaa1100, PubMed: 33237286.

39. Kanehisa, M., Furumichi, M., Tanabe, M., Sato, Y. & Morishima, K. KEGG: new perspectives on genomes, pathways, diseases and drugs. Nucleic Acids Res. 45, D353–D361 (2017). 10.1093/nar/gkw1092, PubMed: 27899662.

40. Kingma, D. P. & Ba, J. A. A method for stochastic optimization. arXiv Preprint ArXiv:1412.6980 (2014). 10.48550/arXiv.1412.6980.

41. Zhao, Q., Zhao, H., Zheng, K. & Wang, J. HyperAttentionDTI: improving drug–protein interaction prediction by sequence-based deep learning with attention mechanism. Bioinformatics 38, 655–662 (2022). 10.1093/bioinformatics/btab715, PubMed: 34664614.

42. Nguyen, N.-Q., Jang, G., Kim, H. & Kang, J. Perceiver CPI: a nested cross-attention network for compound–protein interaction prediction. Bioinformatics 39, btac731 (2023). 10.1093/bioinformatics/btac731, PubMed: 36416124.

43. Kang, H. et al. Fine-tuning of BERT model to accurately predict drug–target interactions. Pharmaceutics 14, 1710 (2022). 10.3390/pharmaceutics14081710, PubMed: 36015336.

44. Singh, R., Sledzieski, S., Bryson, B., Cowen, L. & Berger, B. Contrastive learning in protein language space predicts interactions between drugs and protein targets. Proc. Natl Acad. Sci. U. S. A. 120, e2220778120 (2023). 10.1073/pnas.2220778120, PubMed: 37289807.

45. Rogers, D. & Hahn, M. Extended-connectivity fingerprints. J. Chem. Inf. Model. 50, 742–754 (2010). 10.1021/ci100050t, PubMed: 20426451.

46. Smith, T. F. & Waterman, M. S. Identification of common molecular subsequences. J. Mol. Biol. 147, 195–197 (1981). 10.1016/0022-2836(81)90087-5, PubMed: 7265238.

## References

1 Mead, R. J., Shan, N., Reiser, H. J., Marshall, F. & Shaw, P. J. Amyotrophic lateral sclerosis: a neurodegenerative disorder poised for successful therapeutic translation. Nature Reviews Drug Discovery 22, 185–212 (2023).

